# Centromere clustering stabilizes meiotic homolog pairing

**DOI:** 10.1101/612051

**Authors:** Talia Hatkevich, Vincent Boudreau, Thomas Rubin, Paul S. Maddox, Jean-René Huynh, Jeff Sekelsky

**Affiliations:** Curriculum in Genetics and Molecular Biology, University of North Carolina, Chapel Hill, NC 27599-7264, USA; Department of Biology, University of North Carolina, Chapel Hill, NC 27599-3280, USA; CIRB, Collège de France, PSL Research University, CNRS UMR7241, Inserm U1050, 75005 Paris, France; Integrative Program in Biological and Genome Sciences, University of North Carolina, Chapel Hill, NC 27599-7100, USA

## Abstract

During meiosis, each chromosome must selectively pair and synapse with its own unique homolog to enable crossover formation and subsequent segregation. How homolog pairing is maintained in early meiosis to ensure synapsis occurs exclusively between homologs is unknown. We aimed to further understand this process by utilizing a unique *Drosophila* meiotic mutant, *Mcm5^A7^*. We found that *Mcm5^A7^* mutants are proficient in homolog pairing at meiotic onset yet fail to maintain pairing as meiotic synapsis ensues, causing seemingly-normal synapsis between non-homologous loci. This pairing defect corresponds with a reduction of SMC1-dependent centromere clustering at meiotic onset. Overexpressing SMC1 in this mutant significantly restores centromere clustering, homolog pairing, and crossover formation. These data indicate that the initial meiotic pairing of homologs is not sufficient to yield synapsis between exclusively between homologs and provide a model in which meiotic homolog pairing must be stabilized by SMC1-dependent centromere clustering to ensure proper synapsis.

## INTRODUCTION

Accurate segregation of homologous chromosomes during the first meiotic division is essential to reestablish the diploid genome upon sexual fertilization. To ensure faithful meiosis I chromosomal segregation, homologs must become physically connected in part through crossover formation. To enable homolog crossover events, a series of chromosomal and cellular events occur in early meiotic prophase I (Lake and Hawley 2012) (Figure 1a).

**Figure 1.**
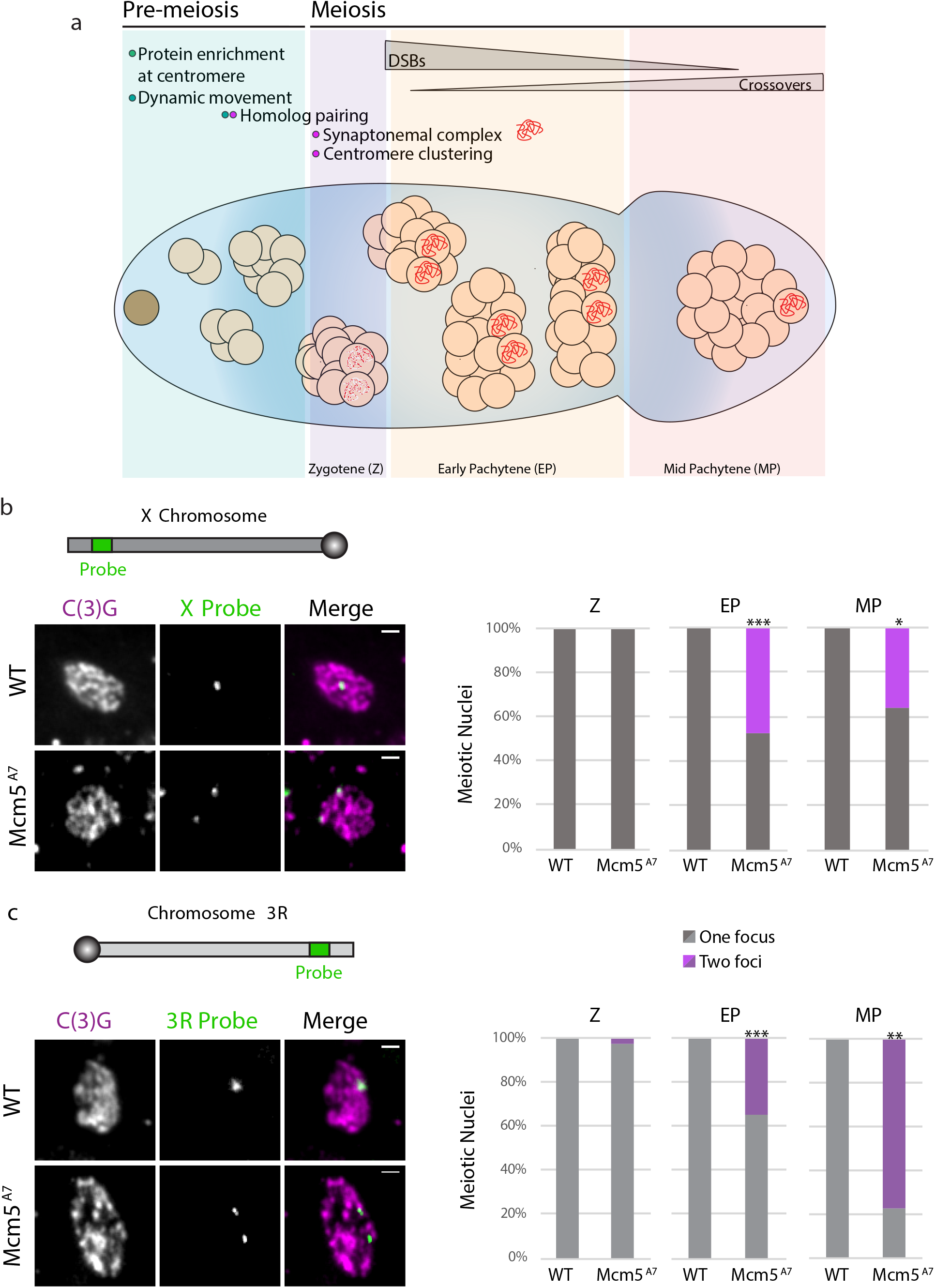
Meiotic pairing is perturbed in *Mcm5^A7^* mutants. a. Schematic depiction of the *Drosophila* germarium. At the anterior portion (the pre-meiotic region, Region 1), the germline stem cell (brown cell) divides to yield a cytoblast, which undergoes four subsequent rounds of division to yield a 16-cell cyst. In the pre-meiotic region, meiotic proteins, such as SMC1 and C(3)G, are enriched at the centromeres, and within the 8-cell cyst, chromosomes exhibit centromere-direct rapid movements. Within the first 16-cell cyst (zygotene; Region 2A), homologous chromosomes pair, centromeres cluster into 1 or 2 groups, and up to four cells initiate meiosis, expressing patches of SC (red dots). As the 16-cell cyst enters early pachytene (EP) (Region 2A), only two continue as pro-oocytes to form full length synaptonemal complex (red). Meiotic double-strand breaks (DSBs) are formed and repaired via homologous recombination (HR) throughout the germarium’s posterior (Regions 2A, 2B) to yield noncrossover and crossover products. At the most posterior tip, signifying mid-pachytene (MP), only one cell within the cyst has been selected to become the oocyte, and all DSBs are repaired. b. Top: schematic of *X* chromosome and relative location of *X*-probe, not drawn to scale. Left: Representative images paired (*WT*) and unpaired (*Mcm5^A7^*) *X*-probes (green) in meiotic cells, indicated by C(3)G expression (magenta). Images are of meiotic nuclei in Region 2A. Scale bar = 1 μm. Right: Quantification of percent paired and unpaired cell in *WT* and *Mcm5^A7^* in Z (*WT n* = 33, *Mcm5^A7^* = 32), EP (WT *n* = 130, *Mcm5A^7^* = 118; \*\*\**p* < 0.0001, chi-square), and MP (*WT n* = 10, *Mcm5^A7^* = 11; *p = 0.01, chi-square). c. Top: schematic of the right arm of chromosome *3* (*3*R) and relative location of *3*R-probe, not drawn to scale. Left: Representative images paired (*WT*) and unpaired (*Mcm5^A7^*) 3R-probes (green) in meiotic cells, represented by C(3)G expression (magenta). *WT* image is of 2A nucleus, *Mcm5^A7^* is of Region 3 nucleus. Right: Quantification of percent paired and unpaired cell in *WT* and *Mcm5A^7^* in Z (*WT n* = 37, *Mcm5A^7^* = 33), EP (*WT n* = 104, *Mcm5^A7^* = 97; \*\*\**p* < 0.0001, chi-square), and MP (*WT n* = 10, *Mcm5A^7^* = 9; \*\**p* = 0.0066, chi-square). Brightness, contrast, and texture (smoothed) of images have been adjusted for clarity.

During or just prior to the onset of meiosis, homologous chromosomes pair along their entire lengths (reviewed in Denise Zickler and Kleckner 2015). Between paired homologs, synapsis, the formation of the synaptonemal complex (SC), ensues. The SC is a tripartite scaffold built between homologs extending the length of the chromosomes and consists of a central region (CR) that is nestled between two lateral elements (LEs), which are successors of cohesin-based chromosome axes formed between sister chromatids. Coincidentally with synapsis, DSBs are formed and repaired using a homologous template via homologous recombination (HR), resulting in crossover formation between homologs (reviewed in Page and Hawley 2004).

Perhaps the most enigmatic event within early meiosis is the mechanism by which a meiotic chromosome selectively pairs and synapses with its unique homologous partner. Initial homolog pairing is believed to be facilitated through early meiotic chromosome movement and telomere or the centromere clustering (for reviews, see Denise Zickler and Kleckner 2015; Alleva and Smolikove 2017; Klutstein and Cooper 2014). However, how homologous pairing is maintained during synapsis to ensure the SC is formed exclusively between homologs is unknown.

The model organism *Drosophila melanogaster* has been used to uncover meiotic mechanisms for over a century (Morgan 1910). In *Drosophila*, prior to meiosis, chromosomes enter the germline unpaired (Figure 1a); throughout the pre-meiotic region, homologous chromosomes gradually pair. In the nuclei at the last mitotic division prior to meiotic onset (in the 8-cell cyst), centromere-directed chromosomal movements occur, presumably ensuring complete homologous pairing (Christophorou et al. 2015; Joyce et al. 2013). Also during pre-meiotic mitotic cycles, meiotic proteins, including the cohesin SMC1, are enriched at the centromere (Khetani and Bickel 2007; Christophorou, Rubin, and Huynh 2013). The onset of meiotic prophase I occurs in the 16-cell cyst. At zygotene, the first cytologically resolved stage of prophase, centromeres are clustered into 1 or 2 groups (Takeo et al. 2011), and the SC nucleates in patches along chromosome arms (Tanneti et al. 2011). As zygotene proceeds into early pachytene, the SC extends between paired chromosomes, yielding full-length SC exclusively between homologs. How these early meiotic events, particularly centromere clustering, contribute to meiotic homologous pairing and synapsis in *Drosophila* is largely unknown.

In this study, we used the *Drosophila* early meiotic program and a unique genetic mutant to investigate how homolog pairing is maintained during meiotic synapsis. We discovered that meiotic homologs in a previously described *Drosophila* mutant, *Mcm5^A7^* (Lake et al. 2007), initially pair, but are unable to maintain pairing during synapsis, suggesting that initial meiotic pairing must be subsequently stabilized by an unknown mechanism to ensure proper synapsis. Using *Mcm5^A7^* as a genetic tool to interrogate pairing stabilization mechanism(s), we show that the meiotic pairing defect and resulting heterosynapsis are due to a lack of SMC1-dependent centromere clustering at meiotic onset. From our results, we suggest a model for proper synapsis in which initial meiotic pairing must be stabilized by centromere clustering, a meiotic event produced by SMC1-enrichment at the centromere and dynamic chromosome movements.

## RESULTS

### Mcm5^A7^ mutants are proficient in initial meiotic pairing but deficient in pairing maintenance

The *Mcm5^A7^* allele, discovered in a meiotic mutant screen, is a missense mutation that changes a conserved aspartic acid residue at the C-terminus, adjacent to the AAA+ ATPase domain. *Mcm5^A7^* mutants have an X-NDJ rate of ~25% that is accompanied with a 90% decrease of crossovers on the X chromosome (Lake et al. 2007). Interestingly, the SC, as shown through staining of the central region (CR) protein C(3)G, appears normal, and DSBs are created and repaired with normal kinetics (Lake et al. 2007). The reason as to why crossovers were severely decreased in *Mcm5^A7^* mutants was unknown at the time of this study.

We hypothesized that a lack of meiotic homolog pairing could result in the severe loss of meiotic crossovers in *Mcm5^A7^* mutants. To test this, we examined the frequencies of X and Chromosome 3R homolog pairing in zygotene, early pachytene, and mid-pachytene meiotic cells using IF/FISH. Zygotene is the earliest cytologically resolved meiotic stage in the *Drosophila* germarium and is defined by the presence of SC patches in the 16-cell cyst. Early pachytene is defined by full-length SC in the early 16-cell cysts (Region 2A of the germarium), and mid-pachytene is defined as the most posterior nucleus in the germarium that expresses full-length SC (Region 3) (Lake and Hawley 2012).

At the X locus, wild-type meiotic cells exhibit one focus throughout zygotene, early pachytene, and mid-pachytene (Z, EP, and MP, respectively, Figure 1b). In *Mcm5^A7^* mutants, we observed one focus at 100% frequency in zygotene. Strikingly, we can resolve two foci in approximately half of the nuclei in *Mcm5^A7^* mutants during early pachytene (\*\*\**p* < 0.0001) and mid-pachytene (\**p* = 0.01, respectively).

Similarly, at the 3R locus wild-type homologous chromosomes are paired at 100% frequency in zygotene, early pachytene, and mid-pachytene (Figure 1c). However, in *Mcm5^A7^* mutants, the homologs of chromosome 3R in zygotene are paired at nearly 100% frequency, yet we can resolve two 3R foci in 35% of early pachytene nuclei (\*\*\**p* < 0.0001) and 78% of mid-pachytene nuclei (\*\**p* = 0.0002).

The above results show that in *Mcm5^A7^* mutants, meiotic chromosomes enter meiosis paired, but as the meiotic nuclei proceed through meiosis, homologous pairing cannot be maintained. This suggests that homolog pairing must be stabilized in early meiosis by an unknown mechanism to ensure accurate synapsis, and in *Mcm5^A7^* mutants, this mechanism is perturbed. Therefore, we reasoned that *Mcm5^A7^* can be used as a genetic tool to interrogate the mechanism that stabilizes meiotic pairing.

### The synaptonemal complex (SC) shows no observable defects in Mcm5^A7^ mutants

Although pachytene homolog pairing is disrupted at a high frequency in *Mcm5^A7^* mutants, the SC, as determined by C(3)G staining, still forms (Lake et al. 2007) (Figure 2a). To explain this, we hypothesize that either (1) the unpaired loci do not correspond with linear SC, or (2) the unpaired loci are forming stable SC with non-homologous loci, creating heterosynapsis. To differentiate between these two, we examined whole mount germaria with IF/FISH and super-resolution microscopy (AIRY Scan) and examined tracts of SC. In wild-type, we can discern that one linear tract of C(3)G is built between the paired X locus (Figure 2b). In *Mcm5^A7^* mutants, separate homologous loci are associated with separate linear tracts of C(3)G, indicating that the unpaired X loci are synapsed with non-homologous loci (see Supplemental Movies 1 and 2). From these data, we conclude that *Mcm5^A7^* mutants have the ability to heterologously synapse.

**Figure 2.**
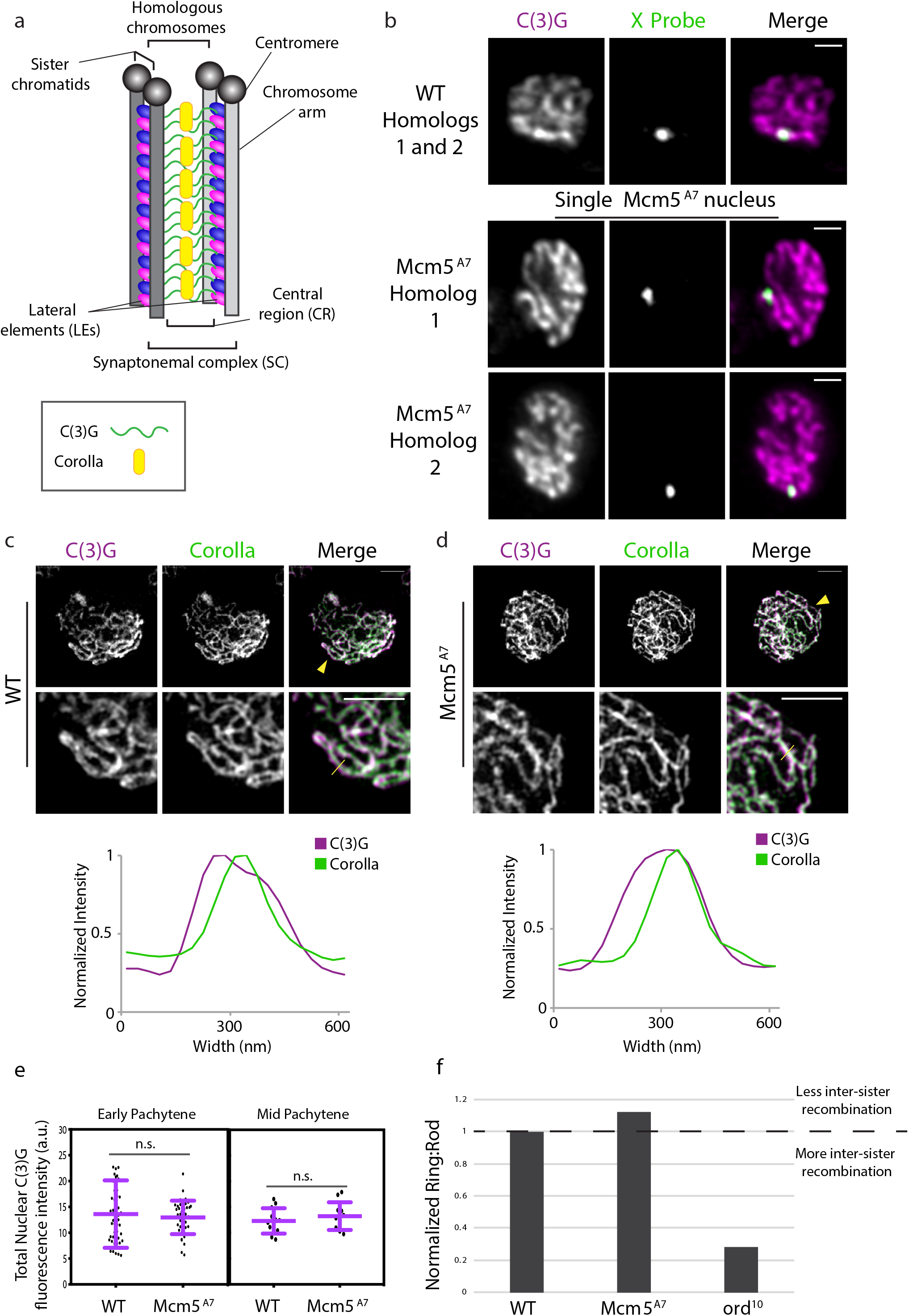
Synaptonemal complex exhibits no observable defects in *Mcm5^A7^* mutants. a. Schematic depiction of SC between two homologous chromosomes. The SC is composed of two lateral elements (LEs) and one central region (CR). The LEs are predecessors of the axial element, which is formed between sister chromatids and are composed of two cohesion complexes (blue and pink ovals. The CR consists, in part, of a C(3)G (green) dimer spanning the LEs, with pillar proteins such as Corolla (yellow) embedded within the CR. Enrichment of proteins at the centromere is not depicted. b. Super-resolution images of C(3)G and *X*-probe in *WT* (paired) and *Mcm5^A7^* (unpaired) in whole-mount germaria. The images of *Mcm5^A7^* is of the same nucleus but of different *Z* slices to capture both *X*-probes. Brightness and contrast have been adjusted for clarity. Scale bar = 1 μm. Refer to Supplemental Movies 1 and 2. c. Top: Representative image of C(3)G (magenta) and Corolla (green) in a *WT* meiotic chromosome spread. Brightness and contrast have been adjusted for clarity. Yellow arrowhead indicates area magnified in lower panel (middle). Scale bar = 2 μm. Middle: Magnification to detail the localization of C(3)G and Corolla. Scale bar = 2 μm. Yellow line indicates the area that was quantified for normalized intensity. Bottom: Normalized intensity of C(3)G and Corolla to demonstrate localization. d. Top: Representative image of C(3)G (magenta) and Corolla (green) in *Mcm5^A7^* meiotic chromosome spread. Yellow arrowhead indicates area magnified in lower panel (middle). Scale bar = 2 μm. Middle: Magnification to detail the localization of C(3)G and Corolla. Scale bar = 2 μm. Yellow line indicates the area that was quantified for normalized intensity. Bottom: Normalized intensity of C(3)G and Corolla to demonstrate localization. e. Left panel: Quantification of nuclear C(3)G signal at early pachytene in *WT* (*n* = 52) and *Mcm5^A7^* (*n* = 41) meiotic nuclei. *p* = 0.5601, unpaired T-test. Data are represented as mean ± SD. Right panel: Quantification of nuclear C(3)G signal in mid-pachytene in *WT* (*n* = 12) and *Mcm5^A7^* (*n* = 11) meiotic nuclei. *p* = 0.3993, unpaired T-test. Data are represented as mean ± SD. Refer to Supplemental Figure 1 for images and further analysis. f. *Mcm5^A7^* (*n* = 1194) and *ord^10^* (*n* = 250) mutants examined for inter-sister recombination through the ratio of Ring chromosome to Rod chromosome transmission. *WT* (*n* = 2574 for *Mcm5^A7^* experiment, *n* = 1204 for *Ord* experiment) was normalized to 1. Ratios above 1 suggest less inter-sister recombination; ratios below 1 suggest more inter-sister recombination. Refer to Table S1 for complete dataset.

To determine the nature of heterosynapsis, we examined the localization of two SC central region (CR) proteins, C(3)G and Corolla (Figure 2a). In wild-type, Corolla co-localizes with C(3)G dimers (Collins et al. 2014), as shown in Figure 2c under structured-illumination microscopy. Under higher resolution, Corolla and C(3)G signal were found to overlap and C(3)G signal is wider, as expected due to its dimer-dimer conformation (Jeffress et al. 2007). In *Mcm5^A7^* mutants, Corolla and C(3)G exhibit a similar localization pattern (Figure 2d). To examine proper CR protein levels, we quantified total C(3)G nuclear signal during early and mid-pachytene in wild-type and *Mcm5^A7^* mutants (Figure 2e). During these timepoints, we see no significant differences between wild-type and *Mcm5^A7^* C(3)G nuclear fluorescence intensity (*p* = 0.5601 and *p* = 0.3993, respectively, unpaired T-test).

The chromosome axis between sister chromatids serves as the lateral element (LE) and provides a barrier to prevent inter-sister recombination (Webber, Howard, and Bickel 2004). To test the function of the LE, we examined inter-sister recombination rates using a genetic ring/rod chromosomal transmission assay (Figure 2f). *Mcm5^A7^* mutants exhibit no decrease in ring:rod transmission (1.1:1 ratio). However, a LE mutant (*ord^10^*) shows a severe decrease in ring:rod transmission to a normalized ratio of 0.28:1. These results suggest that *Mcm5^A7^* mutants exhibit no observable defects in the SC, indicating that seemingly-normal synapsis can occur independent of pairing.

### Centromere-directed chromosome movements are normal in Mcm5^A7^ mutants

We set out to understand how initial pairing of homologs is proficient in *Mcm5^A7^* mutants, despite exhibiting defects in pairing maintenance. Rapid chromosome movements are thought to contribute to homolog pairing (reviewed in Alleva and Smolikove 2017). To determine whether perturbations in centromere-directed chromosome movement contribute to the observed defects in *Mcm5^A7^* mutants, we examined centromere dynamics in 8-cell cysts of wild-type and *Mcm5^A7^* mutant germaria through live cell imaging (Figure 3a, Supplemental Movies 3, 4).

**Figure 3.**
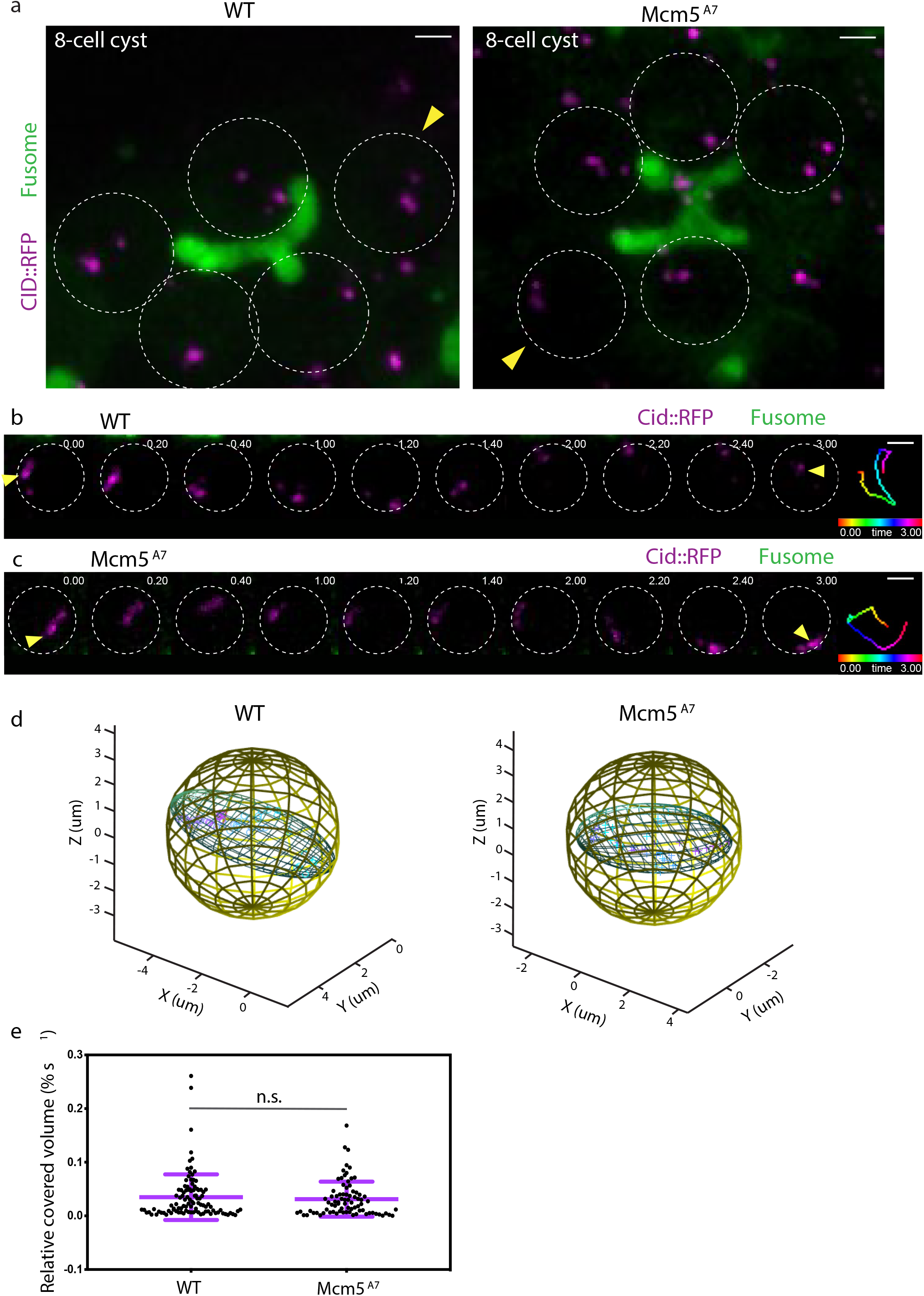
Centromeres in *Mcm5^A7^* mutants exhibit dynamic, rapid movements. a. Projection of Z-sections of live *WT* (left) and *Mcm5^A7^* 8-cell cysts expressing CID::RFP (magenta) and Par-1 ::GFP (fusome, green). Circles represent individual nuclei within the 8-cell cysts. Yellow arrow heads denote representative analyses shown in b. and c. and quantified by time points in d.. Scale bars = 2μm. For videos, refer to Supplemental Video 3 and Video 4. b. Selected projections from one *WT* 8-cell cyst nucleus in (A, indicated by yellow arrow head) over a 3-minute time course. See Video S5 for full movie. c. Selected projections from one *Mcm5^A7^* 8-cell cyst nucleus in (A, indicated by yellow arrow head) over a 3-minute time course. See Video S6 for full movie. Time-colored tracking for CID-RFP dots indicated by yellow arrow heads are shown in right panels for b. and c.. Scale bars = 2μm. d. 3-dimensional representations demonstrating the covered volume of a representative track for all time points in *WT* (50 time points, volume = 12.9 μm^3^) and *Mcm5^A7^* (48 time points, volume = 15.7 μm^3^). e. Distribution of the relative covered volume (raw covered volume/nuclear volume) per second for each track in *WT* (*n* = 103 centromere foci) and *Mcm5A^7^* (*n* = 80 centromere foci). *p* = 0.75, Kolmogorov-Smirnov test. Data are represented as mean ± SD.

In wild-type, a representative centromere track illustrates chromosome movement around the volume of a nucleus (Figure 3b), covering a nuclear volume of 12.9 μm^3^ (Figure 3d, Supplemental Movie 5). A representative centromere track in *Mcm5^A7^* mutants shows similar chromosome movement (Figure 3c), covering a nuclear volume of 15.7 μm^3^ (Figure 3d, Supplemental Movie 6). Of all centromeres analyzed, *Mcm5^A7^* mutants show no significant difference between relative nuclear volume covered compared to wild-type (Figure 3e, *p* = 0.75, Kolmogorov-Smirnov test), demonstrating that *Mcm5^A7^* mutants exhibit centromere-directed chromosome movements similar to wild-type in the 8-cell cyst. Importantly, these data show that centromere-directed chromosome movement may promote initial meiotic homolog pairing but is not sufficient for maintaining homolog pairing.

### Meiotic centromere clustering is defective in Mcm5^A7^ mutants

In *Drosophila*, eight centromeres aggregate into one or two diffraction-limited clusters at the onset of meiosis, which is defined cytologically as zygotene. Centromeres remain clustered through pachytene (Takeo et al. 2011). To determine whether centromere clustering at the onset of meiosis is associated with initial homolog pairing, we quantified the foci number of CID, the CENP-A homolog (Henikoff et al. 2002), in zygotene nuclei in wild-type and *Mcm5^A7^* mutants at zygotene (Figure 4a). We observed a mean of 2 CID foci in wild-type, demonstrating centromere clustering. In *Mcm5^A7^* mutants, we see a significant increase in CID foci, with a mean of 4.8 per nucleus (*p* < 0.001, unpaired T-test). These results show that in *Mcm5^A7^* mutants, centromeres are not heterologously clustered entering meiosis, even though chromosome arms are paired.

**Figure 4.**
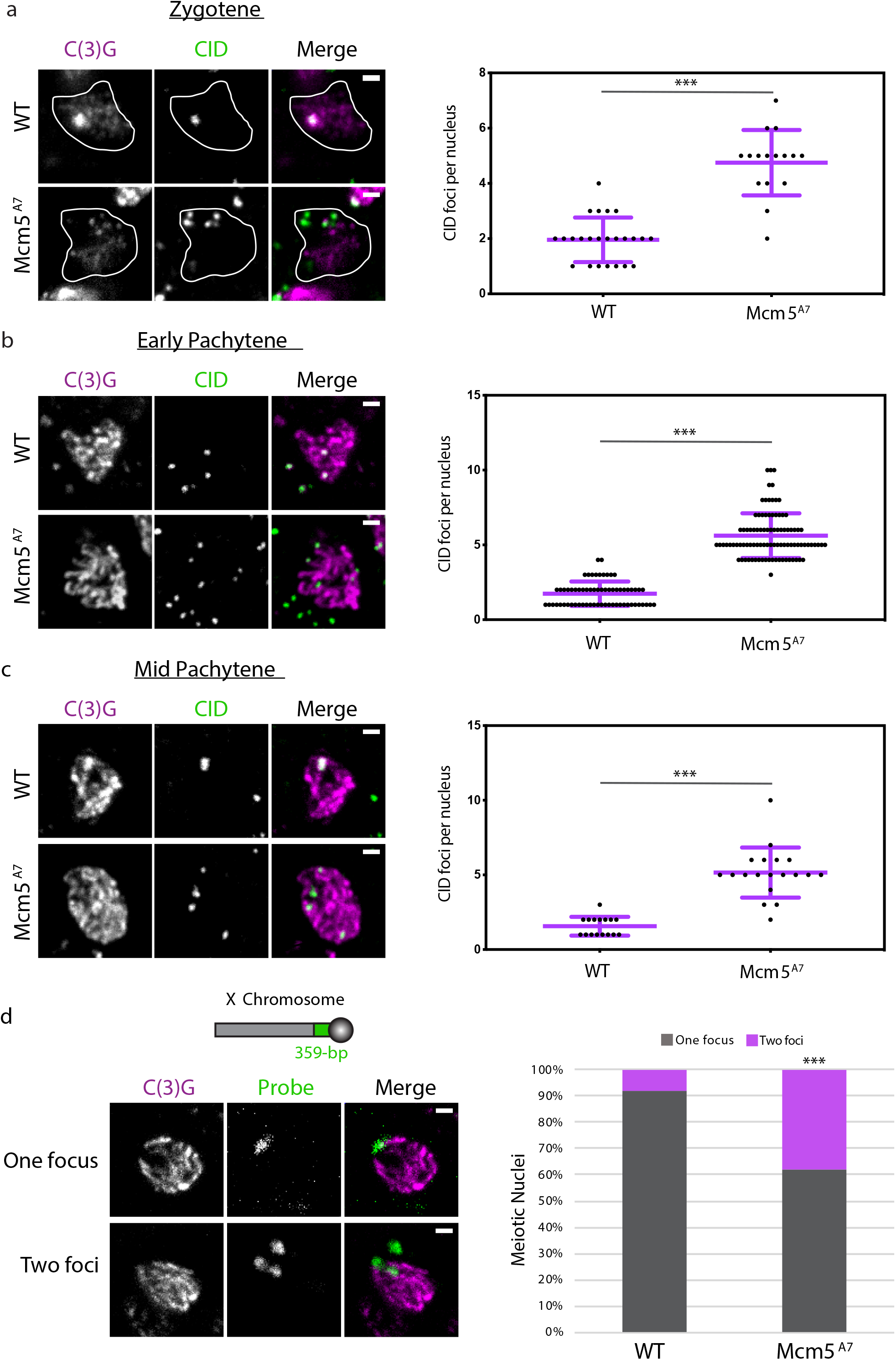
Centromere clustering is disrupted in *Mcm5^A7^* mutants. a. Left: Representative images of centromere clustering, or lack thereof, in wild-type (*WT*) and *Mcm5^A7^* meiotic nuclei located in zygotene. Magenta: C(3)G, green: CID (centromere). In these images, *WT* nucleus contains 1 CID focus, and *Mcm5^A7^* contains 6 CID foci. Scale bar = 1 μm. Circles represent outline of nuclei. CID foci not localized with C(3)G is from adjacent, non-meiotic cells (refer to Supplemental Figure 2a). Right: Quantification of CID foci in zygotene in *WT* (*n* = 24) and *Mcm5^A7^* (*n* = 16). \*\*\**p* < 0.0001, unpaired T-test. Data are represented as mean ± SD. b. Left: Representative images of centromere clustering in *WT* and *Mcm5^A7^* early pachytene nuclei. Magenta: C(3)G, green: CID. In these images, *WT* nucleus contains 2 CID foci, and *Mcm5^A7^* contains 5 CID foci. Scale bar = 1 μm. CID foci not localized with C(3)G is from adjacent, non-meiotic cells (refer to Supplemental Figure 2b). Right: Quantification of early pachytene CID foci in *WT* (*n* = 65) and *Mcm5^A7^* (*n* = 94). \*\*\**p* < 0.0001, unpaired T-test. Data are represented as mean ± SD. c. Left: Representative images of centromere clustering, or lack thereof, in midpachytene *WT* and *Mcm5^A7^* nuclei. Magenta: C(3)G, Green: CID. In these images, *WT* nucleus contains 1 CID focus, and *Mcm5^A7^* contains 4 CID foci. Scale bar = 1 μm. CID foci not localized with C(3)G is from adjacent, non-meiotic cells (refer to Supplemental Figure 2c). Right: Quantification of mid-pachytene CID foci in *WT* (*n* = 16) and *Mcm5^A7^* (*n* = 19). \*\*\**p* < 0.0001, unpaired T-test. Data are represented as mean ± SD. d. Schematic representing relative location of 359-bp locus on Chromosome *X* (not drawn to scale). Top panel: Representative image of meiotic nucleus with 1 359-bp (green) focus (WT, Region 2A). Bottom panel: Representative image of meiotic nucleus with 2 359-bp (green) foci *Mcm5^A7^*, Region 2A). Right: Percentage of nuclei with paired 359-bp loci (one focus) or unpaired (two loci) in *WT* (*n* = 88) and *Mcm5^A7^* (*n* = 63) meiotic nuclei. \*\*\**p* < 0.0001, as determined by two-tailed Fisher’s exact test. Contrast and brightness of all images were adjusted for clarity.

Next, we determined whether centromeres cluster in pachytene in *Mcm5^A7^* mutants. As shown in Figure 4b, we observe a mean of 1.7 CID foci in early pachytene nuclei of wild-type, compared to 5.6 foci in *Mcm5^A7^* mutants (*p* < 0.001, unpaired T-test). In mid-pachytene, *Mcm5^A7^* mutants exhibit a mean of 5.2 CID foci, significantly higher than wild-type (1.6 CID foci; *p* < 0.001, unpaired T-test) (Figure 4c). We conclude that centromere clustering is perturbed throughout early and mid-pachytene in *Mcm5^A7^* mutants.

In the regions assessed, we observed more than four CID foci in most *Mcm5^A7^* nuclei (Figure 4a, b, c), suggesting that homologous centromeres are unpaired. To test this, we examined the pairing frequency of the 359-bp locus, which is adjacent to the *X* centromere (Dernburg, Sedat, and Hawley 1996) (Figure 4d). As previously reported, this locus is paired in ~90% of meiotic cells (Joyce et al. 2013); however, in *Mcm5^A7^* meiotic nuclei, 359-bp locus pairing is significantly reduced to 61% (*p* < 0.001, two-tailed Fisher’s exact test). From these results, we conclude that meiotic homologous centromere pairing and heterologous centromere clustering are severely decreased, if not eliminated, in *Mcm5^A7^* mutants. These data indicate that a decrease in meiotic centromere clustering is associated with defects in homologous chromosome pairing but not pre-meiotic pairing. Also, these results suggest that mechanisms regulating chromosome arm pairing and chromosome centromere pairing may be distinct.

### SMC1 localization is reduced specifically at the centromere in Mcm5^A7^ mutants

Centromere clustering is perturbed in sine cohesin and SC mutants (Takeo et al. 2011; Christophorou, Rubin, and Huynh 2013; Tanneti et al. 2011), suggesting that specific proteins at the centromeres are required for the aggregation of centromeres. Since we see no decrease of C(3)G at centromeres in *Mcm5^A7^* mutants compared to wild-type (Supplemental Figure 1), we hypothesized that a lack of centromeric cohesion in meiosis may contribute to the decrease in centromere clustering. To test this, we investigated chromosome associatied-SMC1 using meiotic chromosome spreading (Khetani and Bickel 2007) (Figure 5a). In wild-type meiotic nuclei, SMC1 is enriched at the centromere (green arrowhead); because SMC1 contributes to the axial element (AE) formed between sister chromatids, which later serves as the LE of the SC, SMC1 is visualized at the arm as thread-like (yellow arrowhead, dotted line). In *Mcm5^A7^*, SMC1 exhibits thread-like patterning along the arms, but SMC1 enrichment at the centromere appears to be compromised.

**Figure 5.**
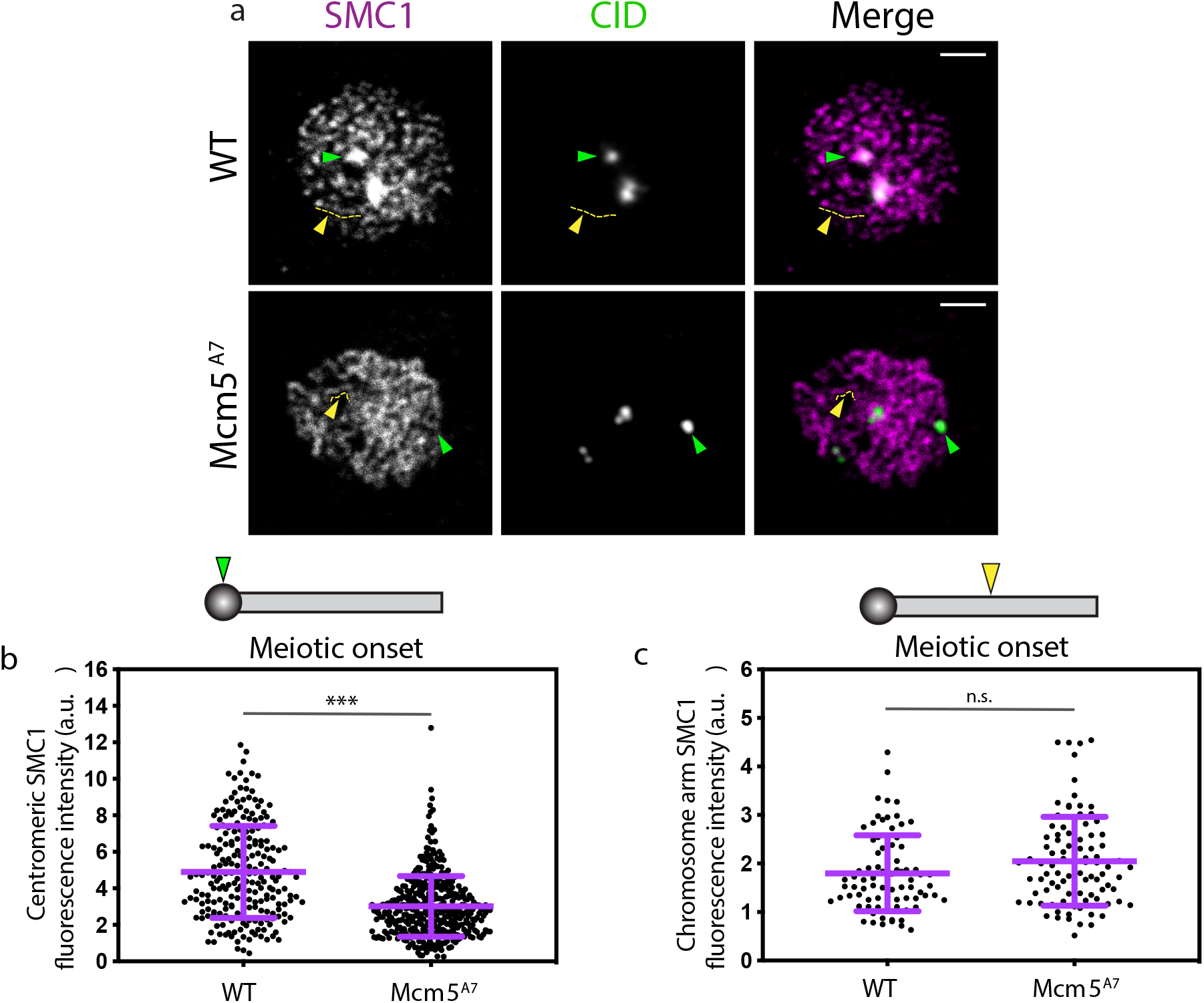
Centromeric SMC1 is significantly reduced in *Mcm5^A7^* mutants. a. Representative images of chromosome spreads in *WT* and *Mcm5^A7^* meiotic nuclei examining localization of SMC1 (magenta) and CID (green). Green arrow: SMC1 enrichment at the centromere, yellow arrow and tract: SMC1 along the chromosome arm. Scale bar = 2 μm. Contrast and brightness of images were adjusted for clarity. b. Quantification of SMC1 at CID foci in meiotic nuclei at meiotic onset (zygotene + early pachytene, Region 2A) at *WT* (*n* = 225) and *Mcm5^A7^* (*n* = 398) meiotic centromeres. \*\*\**p* < 0.0001, unpaired T-test. Data are represented as mean ± SD. c. Quantification of SMC1 at chromosome arm in meiotic nuclei at meiotic onset (zygotene + early pachytene, Region 2A) in *WT* (*n* = 81) and *Mcm5A^7^* (*n* = 93) meiotic nuclei. *n.s*. = 0.0548, unpaired T-test. Data are represented as mean ± SD. Refer to Supplemental Figure 3 for representative images.

We quantified SMC1 localization at the centromere and along the arms at meiotic onset (defined cytologically as zygotene and early pachytene, which cannot be distinguished based on SMC1 patterning), when we hypothesize SMC1 enrichment would be essential for centromere clustering. Strikingly, at meiotic onset, SMC1 is significantly reduced in *Mcm5^A7^* mutants at the centromere, but not along the arms (\*\*\**p* < 0.001 and *p* = 0.0548, respectively, Figures 5b, 5c). These data indicate that SMC1 enrichment specifically at the centromere is perturbed in *Mcm5^A7^* mutants during meiotic onset when non-homologous centromeres should cluster.

### Increasing centromere clustering ameliorates pairing defects

Using *Mcm5^A7^* mutants, we observed that a decrease in centromeric SMC1 at meiotic onset is associated with a reduction in meiotic centromere clustering and homologous chromosome pairing in pachytene, but not chromosome pairing in zygotene. Thus, we hypothesized that the centromeric-SMC1 defect at meiotic onset causes the reduction in centromere clustering, and that centromere clustering defects cause the defect in pairing maintenance.

To test this hypothesis, we attempted to restore SMC1 localization at the meiotic centromere in *Mcm5^A7^* mutants by exogenously expressing SMC1 (Gyuricza et al. 2016) in the background of *Mcm5^A7^* (*nos>Smc1; Mcm5^A7^*) (Supplemental Figure 5). Using quantitative microscopy, we found that centromeric-SMC1 is significantly higher in *nos>Smc1; Mcm5^A7^* than in *Mcm5^A7^* mutants at meiotic onset (\*\*\**p* < 0.0001, unpaired T-test) (Figure 6a). We next assayed centromere clustering at early pachytene, when we first observe pairing defects in *Mcm5^A7^* mutants (Figure 1b); as shown in Figure 6b, centromere clustering was significantly increased in *nos>Smc1; Mcm5^A7^* as compared to *Mcm5^A7^* (\*\*\**p* < 0.0001, unpaired T-test), indicating that the increase in centromeric-SMC1 localization at meiotic onset partially rescues the early pachytene centromere clustering deficiency in *Mcm5^A7^* mutants.

**Figure 6.**
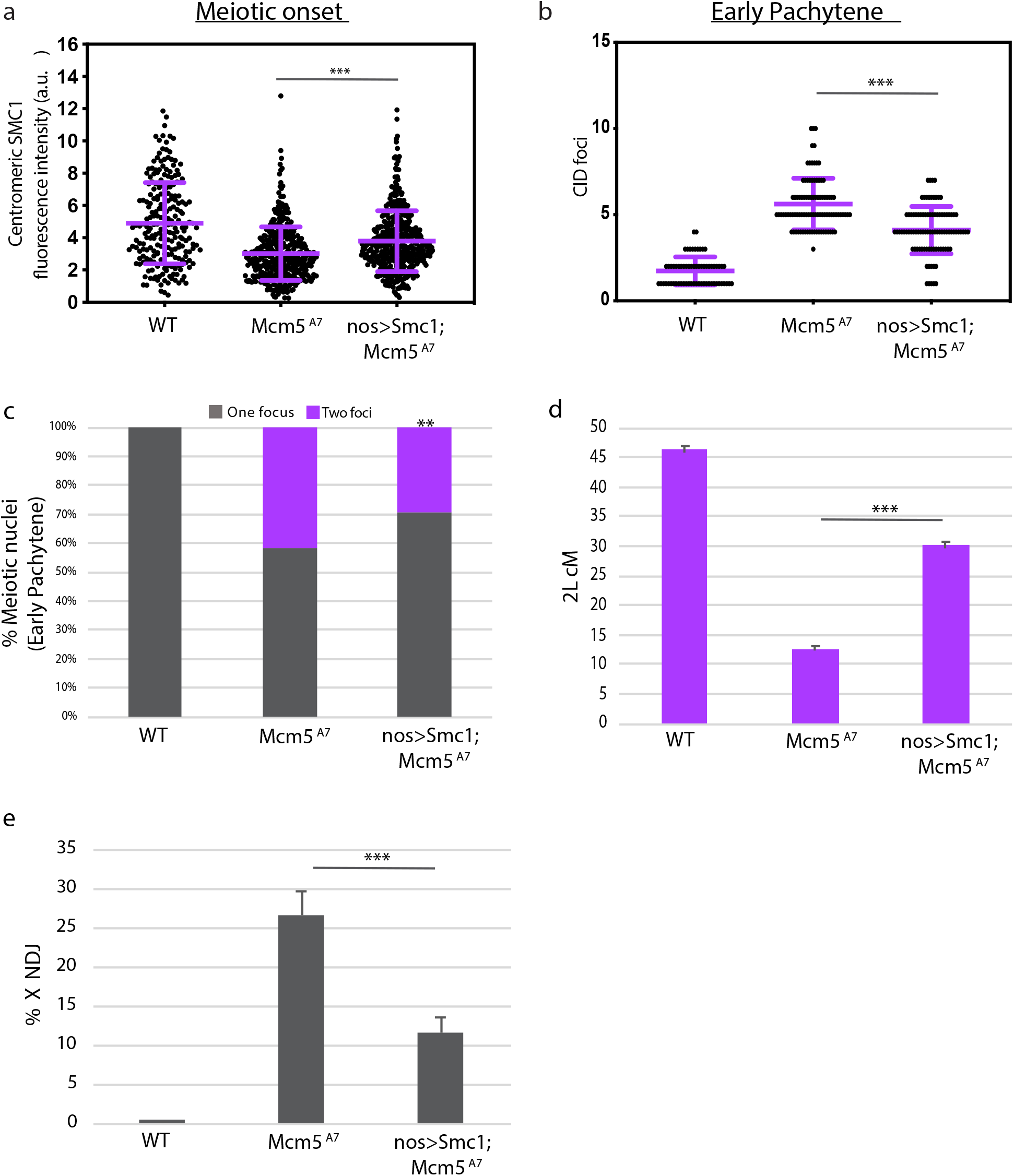
Overexpression of SMC1 in *Mcm5^A7^* mutants rescue clustering, pairing, crossover formation, and NDJ. a. Quantification of SMC1 signal at the centromeres at *WT, Mcm5^A7^*, and *nos>Smc1; Mcm5^A7^* (*n* = 427) meiotic centromeres at meiotic onset. See Supplemental Figure 4 for representative images of *nos>Smc1; Mcm5^A7^* nuclei. *WT* and *Mcm5^A7^* data are repeated from Figure 5. \*\*\**p* < 0.0001, unpaired T-test. Data are represented as mean ± SD. b. Number of centromeres (CID foci) in *WT, Mcm5^A7^*, and *nos>Smc1; Mcm5^A7^* (*n* = 94) meiotic nuclei at early pachytene. *WT* and *Mcm5^A7^* data are repeated from Figure 2. \*\*\**p* < 0.0001, unpaired T-test. Data are represented as mean ± SD. c. Percent of total paired and unpaired in *WT, Mcm5^A7^*, and *nos>Smc1; Mcm5^A7^* (total *n* = 169) nuclei at early pachytene, combining *X*-probe and 3R-probe data. *WT* and *Mcm5^A7^* data are repeated from Figure 4 and are represented as *X*-probe plus 3R-probe early pachytene data. Significance comparing *Mcm5^A7^* and *nos>Smc1, Mcm5^A7^*: \*\**p* = 0.0002, chi-square d. Crossover levels on chromosome 2L as shown in cM in *WT* (*n* = 4222)(Hatkevich et al. 2017), *Mcm5^A7^* (*n* = 2070), and *nos>Smc1; Mcm5^A7^* (*n* = 933). \*\*\**p* < 0.001, chi-square. Data are represented as mean ± 95% CI. Refer to Table S2 for full 2L crossover dataset. e. NDJ of the *X* chromosome in *WT* (0.07%, *n*= 3034), *Mcm5^A7^* (26.5%, *n* = 1979), *nos>Smc1; Mcm5^A7^* (11.5%, *n* = 2282). \*\*\**p* < 0.0001 (Zeng et al. 2010). Data are represented as mean ± 95% CI. Refer to Table S3 for full NDJ dataset.

We reasoned that if SMC1-dependent centromere clustering is partially rescued at early pachytene in *nos>Smc1; Mcm5^A7^*, then the pairing defect at this stage will be attenuated. To test this, we examined pairing frequency of *X* and 3R at early pachytene in *nos>Smc1; Mcm5^A7^* flies (Figure 6c). We see a significant pairing increase in *nos>Smc1; Mcm5^A7^* mutants compared to *Mcm5^A7^* mutants (pairing frequency of 71% and 59%, respectively, \*\**p* = 0.0066, chi-square). From these data, we propose that SMC1-dependent centromere clustering in early meiosis promotes the stabilization of meiotic homolog pairing, giving rise to homosynapsis.

We initially hypothesized that a lack of homolog pairing results in the loss of meiotic crossovers in *Mcm5^A7^* mutants. We reasoned that in the presence of heterosynapsis, as seen in *Mcm5^A7^* mutants, meiotic DSBs cannot be repaired into crossovers because no homologous template is available (for a review on homologous recombination, see Hunter 2015). To test this hypothesis, we measured crossovers across chromosome 2L in wild-type, *Mcm5^A7^*, and *nos>Smc1; Mcm5^A7^* mutants (Figure 6d). Wild-type flies exhibit a crossover level of 45.8 cM, while the *Mcm5^A7^* crossover level is significantly decreased to 12.3 cM (\*\*\**p* < 0.0001, chi-square). In *nos>Smc1; Mcm5^A7^* mutants, crossover level is significantly increased to 29.8 cM (\*\*\**p* < 0.0001, chi-square, as compared to *Mcm5^A7^* mutants). These results indicate that the pairing defect and heterosynapsis during early pachytene is, at least in part, the cause for the loss of crossovers in *Mcm5^A7^* mutants.

Because crossover level is partially rescued in *nos>Smc1; Mcm5^A7^* flies, then the high nondisjunction rate in *Mcm5^A7^* should be lessened when SMC1 is overexpressed in these mutants. We observe that *nos>Smc1; Mcm5^A7^* mutants have a significant decrease in X-NDJ as compared to *Mcm5^A7^* mutants (NDJ rate of 11.5% and 26.5%, respectively, Figure 6e) (***p<0.0001). Overall, these studies show that germline overexpression of SMC1 can restore SMC1 at the centromere in *Mcm5^A7^* mutants in early pachytene, leading to increases in centromere clustering, homolog pairing (and homosynapsis), crossover formation, and a decrease in NDJ.

## DISCUSSION

For a successful meiosis, homolog pairing must be maintained during synapsis, but how homologs remain paired as synapsis ensues is unclear. At the beginning of this study, we hypothesized that the crossover defect in *Mcm5^A7^* mutants was due to a homolog pairing deficiency. Our FISH results support this hypothesis (Figure 1) and revealed that homolog pairing is reversible, and if not stabilized, can cause seemingly-normal heterosynapsis (Figure 2). Centromere-directed chromosome movements occur in *Mcm5^A7^* mutants (Figure 3), presumably yielding initial chromosome arm pairing; however, centromere clustering is perturbed (Figure 4). SMC1 enrichment at the centromere is decreased in *Mcm5^A7^* mutants (Figure 5), and an increase in centromeric SMC1 rescues this deficiency and downstream meiotic defects, including centromere clustering, pairing, crossover formation, and segregation (Figure 6). From our data, we propose that meiotic centromere clustering stabilizes initial homolog pairing to give rise to secure meiotic pairing and homosynapsis (Figure 7).

**Figure 7.**
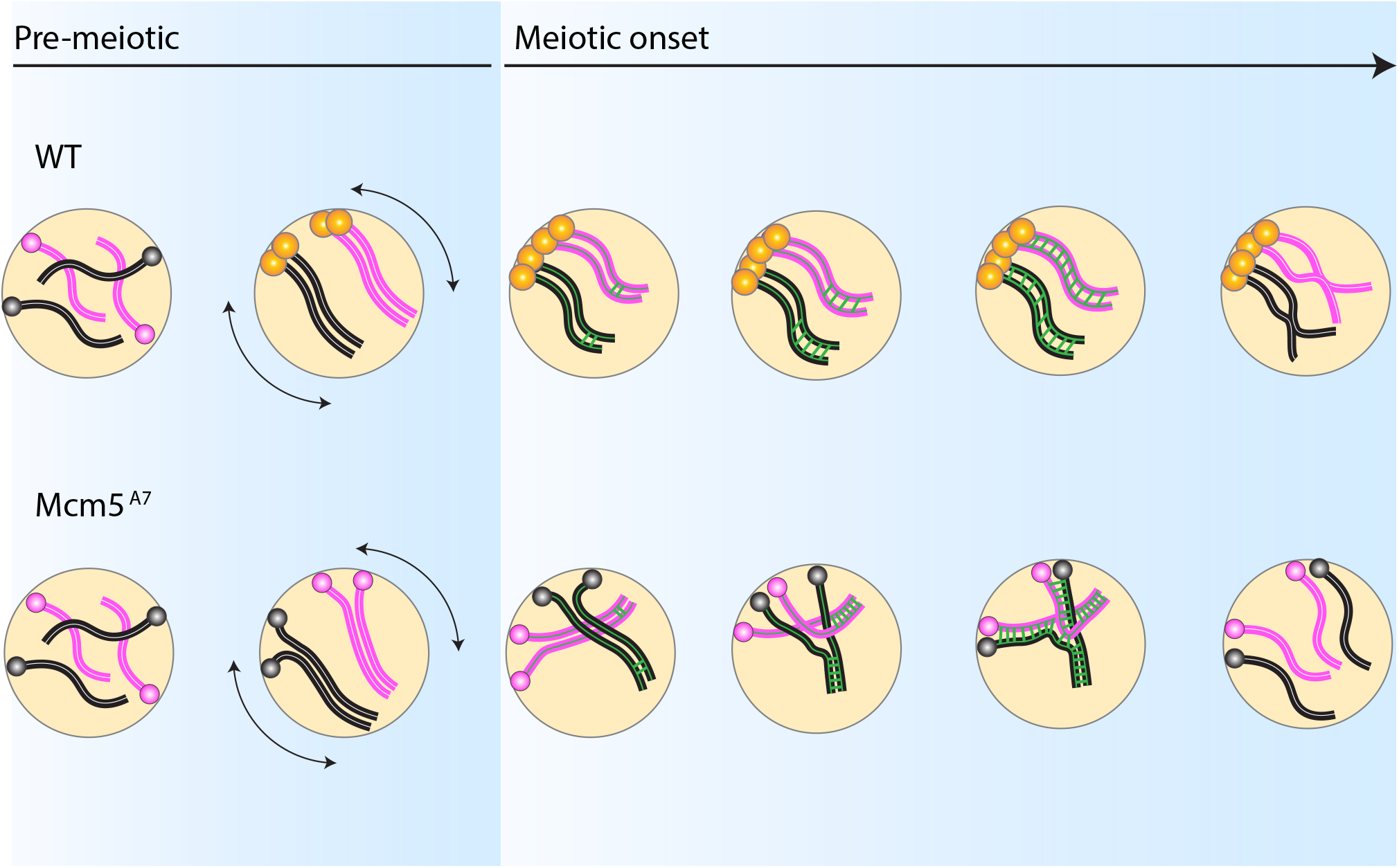
Centromere clustering-dependent pairing model. *WT*: In pre-meiotic cysts, homologous chromosomes (pink = homolog pair 1, black = homolog pair 2) enter the germline unpaired. During pre-meiotic cell cycles, chromosome arms and centromeres pair, with centromeres anchored at the nuclear envelope. Prior to meiotic onset, SMC1 is enriched at the centromeres (yellow) and centromere-directed chromosome movement (double-headed arrows) occurs. These events yield centromere clustering at meiotic initiation. As synapsis nucleates along arms (green bars), paired chromosomes are able to withstand opposing forces because of physical stabilization provided by centromere clustering, permitting homosynapsis. After DSB formation and repair (not depicted), crossovers between homologs are formed, promoting proper disjunction at the end of meiosis I. *Mcm5^A7^*: Chromosomes enter the germline unpaired, and centromeres are attached to the nuclear envelope. In pre-meiotic cycles, chromosomes initially pair, but centromeres do not. Centromere-directed chromosome movements occur, but SMC1 is not enriched at the centromere, causing a lack of centromere clustering at meiotic onset. As the SC nucleates at the arms, opposing forces push the paired chromosome arms apart. Synapsis spreads between the nearest chromosomal regions, independent of homology, yielding high frequency of heterosynapsis. During heterosynapsis, DSBs are not repaired by HR, yielding non-recombinant chromosomes that nondisjoin at the end of Meiosis I.

### A centromere clustering-dependent homolog pairing model in Drosophila

Prior to meiosis, cellular events occur to prepare chromosomes for meiotic pairing and synapsis. Meiotic cohesins are loaded onto centromeres (Khetani and Bickel 2007), and homologous chromosomes pair (Joyce et al. 2013; Christophorou, Rubin, and Huynh 2013), partly due to centromere-directed movements in the division prior to meiotic onset (Christophorou et al. 2015). We propose a model in which initial chromosomal pairing is stabilized throughout early meiosis by SMC1-dependent centromere clustering (Figure 7).

According to this model, the enrichment of SMC1 at the centromere and chromosome movements in pre-meiotic stages yield centromere clustering at meiotic onset. While chromosome arms and centromeres enter meiosis paired, heterologous centromere clustering serves as a mechanism to stabilize the pairing, resisting forces generated by synapsis nucleation and/or diffusion that may otherwise push paired chromosomes apart. As the SC extends between the arms of homologs, DSBs are formed and subsequently repaired via HR to yield crossovers, which promote accurate disjunction at the end of meiosis.

In *Mcm5^A7^* mutants, coordinated pre-meiotic centromere-directed movements occur, yet there is not sufficient SMC1 enriched at the centromere to yield centromere clustering. Thus, at meiotic onset, arms are paired, but centromeres are not clustered. As SC nucleation occurs, the stabilization provided by centromere clustering is absent and chromosome arms move freely in response to SC nucleation and/or diffusion. As synapsis extends, the SC is formed between nearby chromosomes, regardless homology, yielding heterologous synapsis. During instances of heterosynapsis, DSBs are made but cannot be repaired via HR without a homologous template. Therefore, overall crossover levels are reduced, and nondisjunction occurs at high frequency in *Mcm5^A7^* mutants.

The centromere clustering-dependent pairing model highlights that initial meiotic pairing is not sufficient to yield homosynapsis, indicating that pairing may be a two-step process. Initial homolog pairing must occur, but a stabilization step must be enforced for proper synapsis. In *Drosophila*, this stabilization is provided by SMC1-dependent centromere clustering. We propose that, to ensure stabilization of the initial pairing event, centromere clusters act as anchors at the nuclear envelope, maintaining the rigid AE (which runs along the entire length of the arm to the centromere) of each chromosome in proximity of its homolog.

Although meiotic pairing programs vary among organisms, we suggest that the centromere clustering-dependent pairing model can be universally applied. In *Drosophila* and *C. elegans*, meiotic pairing is independent of meiotic recombination. In contrast, meiotic pairing in organisms such as yeast, plants, and mice require DSB formation (although recombination-independent alignment is required for pairing in these organisms)(Denise Zickler and Kleckner 2015). In DSB-dependent pairing programs, homologs are considered paired at ~400 nm, where DSB-mediated interhomolog interactions can be visualized as bridges (Albini and Jones 1987). However, contemporaneous with DSB formation, centromeres are coupled or clustered (reviewed in Da Ines and White 2015). We speculate that these centromere interactions stabilize the DSB-dependent arm pairing to ensure synapsis exclusively between homologs.

### Pairing and subsequent synapsis in Drosophila

This study reveals the interesting phenomenon of extensive, stable heterosynapsis. Extensive heterosynapsis has been previously reported in *C. elegans* (Sato-Carlton et al. 2014.; Couteau et al. 2004; Couteau and Zetka 2005; Martinez-Perez and Villeneuve 2005) and yeast (Zickler and Kleckner 1999) with variable SC defects. Though we cannot rule out SC aberrations in *Mcm5^A7^* mutants, our data reveal no structural defects, supporting the notion that “normal” synapsis is largely homology-independent(Rog and Dernburg 2013). However, results from this study suggest that synapsis initiation may require homology.

In *Drosophila*, synapsis initiates at the arms in patches during zygotene (Tanneti et al. 2011). In *Mcm5^A7^* mutants, synapsis initiation between paired homologs appears normal in zygotene; rather, the elongation of SC from the presumed homologous initiation patches fails to occur between homologs. Therefore, it appears that the initiation of synapsis requires homology, unlike SC elongation. Similar to what has been observed in other organisms (reviewed in Rog and Dernburg 2013), we speculate that synapsis elongation is processive, such that once nucleated, the SC central region will build between two non-homologous chromosome axes that are in close proximity. Future studies determining the degree of heterosynapsis along entire chromosome arms in *Mcm5^A7^* mutants may provide more insight into how synapsis and homology interact in flies.

The heterosynapsis observed in this study also negates the long-standing assumption in *Drosophila* that stable synapsis occurs only between homologs, *i.e.*, if synapsis occurs in a mutant, then the mutant is proficient in pairing. Thus, mutants in *Drosophila* (and perhaps in other organisms) that have been previously believed to be competent in pairing due to the presence of stable SC should be revisited and tested for pairing deficiencies. Doing so could result in novel pairing mutants and aid in further understanding of how a meiotic chromosome pairs and synapses with its unique homolog.

## MATERIALS and METHODS

### Experimental model details

In all experiments, *Drosophila melanogaster* adult females 3-10 days old were used. Flies were maintained on standard medium at 25°C. *Drosophila* nomenclature used in this study was generalized for readership. Nomenclature and specific genotypes are listed below.

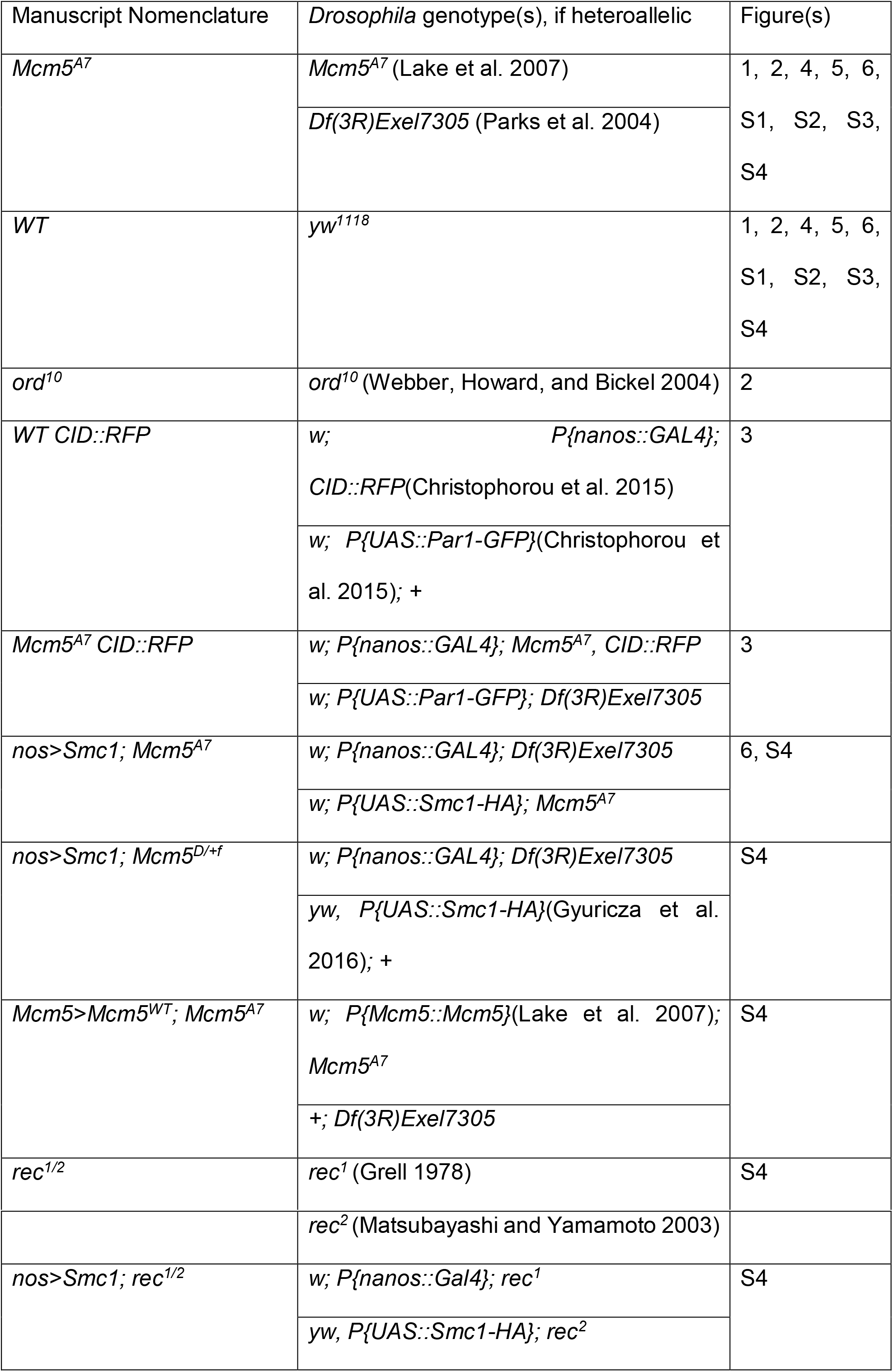

### Experimental details

#### Genetic assays

X chromosome NDJ was evaluated by scoring the progeny from virgin females of desired genotype crossed with *y cv v f / T(1:Y)B^S^* males. Viable exceptional *XXY* females have *Bar* eyes, and viable exceptional X0 males have *Bar*^+^ eyes and are *y cv v f*. To adjust for inviable exceptional males and females, viable exceptional class was multiplied by 2. % *X*-NDJ = 100* ([2*viable exceptional females] + [2*viable exceptional males])/total progeny. Statistical comparisons were performed as in Zeng *et al.*, 2010.

Crossovers on chromosome 2L were measured by crossing virgin *net dpp^ho^ dp b pr cn* / + females of desired genotype to *net dpp^ho^ dp b pr cn* males. Vials of flies were flipped after three days of mating. Resulting progeny were scored for all phenotypic markers. Similarly, crossovers on chromosome *X* were measured by crossing virgin *y sc cv v g f y*^+^ / + females to *y sc cv v g f* males. Progeny were assessed for all phenotypic markers.

To calculate intersister recombination, *R(1)2, y^1^ w^hd80k17^ f^1^/y^1^* females with desired genotype were crossed to *y^1^ w^1118^* and progeny was scored for phenotypic markers. Exceptional progeny were able to be distinguished through phenotypes and rates were adjusted to reflect only normal progeny, as in Webber, Howard and Bickel, 2004.

##### Dissection and immunofluorescence (IF) of whole mount germaria

Ten three-to five-day old virgin females of desired genotype were fattened overnight with yeast paste in vials with ~5 males of any genotype. Ovaries were dissected in fresh 1x PBS and incubated in fixative buffer for 20 minutes. Fixative buffer: 165 μL of fresh 1x PBS, 10 μL of N-P40, 600 μL of heptane, and 25 μL of 16% formaldehyde. After being washed 3 times in 1x PBS + 0.1% Tween-20 (PBST), ovaries were incubated for 1 hour in 1 mL PBST + 1% BSA (10 mL of PBST + 0.1 g BSA). Ovaries were incubated overnight in 500 μL primary antibody diluted in 1 mL PBST + 1% BSA at 4° C on rocking nutator. After being washed 3x in PBST, ovaries were incubated in 500 μL secondary antibody diluted at 1:500 in PBST + 1% BSA for 2 hours under foil. Ovaries were mounted in 35 μL of ProLong Gold + DAPI on microscope slide using fine-tip forceps to spread ovaries.

Antibodies for C(3)G(Anderson et al. 2005), SMC1(Khetani and Bickel 2007), and CID (Active Motif) were used. For Figures 4a, 4b, 4c, S2a, S2b, S2c: Images of whole mount germaria were taken Zeiss LSM880 confocal laser scanning microscope using 63x/0.65 NA oil immersion objective, with a 2x zoom using ZEN software. Images were saved as .czi files and processed using FIJI(Schindelin et al. 2012). For Figures S1a, S1d, S3a, S4a: Images were taken on Nikon A1R point-scanning confocal microscope using 60x 1.49 NA oil immersion objective. Images were saved as .nd2 files and quantified as described below.

#### Dissection and IF of chromosome spreads

Before dissection, 25 mL of fixative, 5 mL of hypo-extraction buffer, and 500 μL of 100 mM sucrose were prepared. Fixative (25 mL): 23.0875 mL water, 1.5625 mL 16% formaldehyde, at 350 μL of 10% Triton-X (1 mL of Triton-X + 9 mL water). Hypo-extraction buffer (5 mL): 3.685 mL water, 250 μL 600 mM Tris (pH 8.2), 500 μL 170 mM Trisodium Citrate Dihydrate, 50 μL 500 mM EDTA, 2.5 μL 1.0 M DTT, 12.5 μL 200 mM Pefabloc (hypo-extraction buffer is good for only 2 hours). 100 mM Sucrose (500 μL): 100 μL 500 mM sucrose + 400 μL water).

Ovaries were dissected in 1x PBS and rinsed once in hypo-extraction buffer. Ovaries were incubated for 20 minutes in hypoextraction buffer and transferred to sucrose and minced. A super-frost slide was dipped into the fixative for 15 seconds. 10 μL of minced ovary tips were transferred onto the middle edge of the long side of the slide and rolled to allow spreading. Slides were dried very slowly overnight in a closed humidified chamber. Once dried, slides were incubated with 500 μL of blocking (5% normal goat serum (NGS), 2% BSA, 0.1% Triton-X in 1x PBS). Slides were rinsed 3 times in B-PBSTx (0.1% BSA, 0.1% Triton-X in 1x PBS). 250 μL of primary antibodies diluted in B-PBSTx were incubated under parafilm overnight in humidifying chamber. Slides were washed 3 times with PBSTx (0.1% Triton-X in 1x PBS). Secondary antibodies were diluted at 1:400 in B-PBSTx. 100 μL of diluted secondary were added onto slide under parafilm and incubated for an hour. Slides were rinsed 3 times in PBSTx and washed three time for 10 minutes in PBSTx in Coplin jar. Slides were incubated swith 400 μL DAPI (1 ug/ml) in 1x PBS for 10 minutes in dark and washed in 1x PBS. Coverslips were mounted with ProLong Gold.

Antibodies for C(3)G(Anderson et al. 2005), Corolla(Collins et al. 2014), SMC1(Khetani and Bickel 2007), and CID (Active Motif) were used. Figure 5a: Images were taken on Zeiss LSM880 confocal laser scanning microscope using 63x/0.65 NA oil immersion objective with a 2x zoom using ZEN software. Images were saved as .czi files and processed using FIJI (Schindelin et al. 2012). Figures 2c and 2d: Images were taken on Nikon N-SIM using Elements software. Images were saved as .nd2 files and processed using FIJI(Schindelin et al. 2012).

#### Generation of fluorescence in situ hybridization (FISH) probes

DNA from desired BAC clones (BAC PAC RPCI-98 Library) was extracted from MIDI-prep culture. For X probe (Figure 1b), six BAC clones were used, spanning cytological bands 6E-7B. Clones: 17 C09, 06 J12, 35 J16, 20 K01, 35 A18, 26 L11. For 3R probe (Figure 1c), six BAC clones were used, spanning cytological bands 93A-93E. Clones: 19 P12, 05 I01, 20 N14, 10 M16, 06 L13, 34 E13. The BAC-derived template DNA was used in a nick-translation reaction to generate euchromatic biotinylated DNA probes, as described below.

For one BAC clone DNA template, the following was added into a 0.5 mL tube: 5 μL 10X DNA Pol I buffer, 2.5 μL dNTP mix (1 mM each of dCTP, dATP, dGTP), 2.5 μL biotin-11-dUTP (1 mM), 5.0 μL 100 mM BME, 10 μL of freshly diluted dDNase I, 1 μL DNA Pol I, 1 ug of template DNA, water up to 50 μL. Reaction was incubated at 15° C in thermocycler for 4 hours and eluted in 20 μL TE. Concentration was determined using Qubit kit and diluted to a final concentration at 2 ng/μL in hybridization buffer. Hybridization buffer: 2x Saline-Sodium Citrate (SSC) buffer, 50%formamide, 10% w/v dextran sulfate, 0.8 mg/mL salmon sperm DNA.

The 359-bp probe (Figure 4d) was ordered from Integrative DNA Technologies (IDT, www.idtdna.com) with 5’ Cy5, resuspended in 1x TE at 100 μM. Sequence for 359-bp probe (5’ to 3’): Cy5-GGGATCGTTAGCACTGGTAATTAGCTGC.

#### FISH/IF of whole mount germaria

Ovaries were dissected in fresh 1x PBS and incubated in fixative buffer for 4 minutes. Fixative buffer: 100 mM sodium cacodylate (pH 7.2), 100 mM sucrose, 40 mM potassium acetate, 10 mM sodium acetate, 10 mM EGTA, 5% formaldehyde. Ovaries were transferred to 0.5 ml tube filled with 2x SSCT (5 ml 20x SSC, 50 μL Tween, 45 mL water) and washed four times in 2x SSCT, 3 minutes each. Ovaries were washed 10 minutes in 2x SSCT + 20% formamide, 10 minutes 2x SSCT + 40% formamide, and then two times for 10 minutes each in 2x SSCT + 50% formamide. Ovaries were incubated at 37° C for 4 hours, at 92° C for 3 minutes and then 60° C for 20 minutes. Ovaries were transferred to tube with 36 μL of BAC-generated probe (diluted in hybridization buffer) or with 35 μL of hybridization buffer and 1 μL of IDT-generated probe. Ovaries were incubated in the thermocycler for 3 minutes at 91° C then at 37° C overnight and then washed with 2x SSCT + 50% formamide for 1 hour at 37° C. Ovaries were washed in 2x SSCT + 20% formamide for 10 minutes at room temperature and rinsed in 2x SSCT four times quickly. Ovaries were incubated for 4 hours in blocking solution (6 mg/mL NGS in 2x SSCT) and then washed three times quickly in 2x SSCT. Ovaries were incubated overnight in primary antibody diluted in 2x SSCT at room temperature. Ovaries were washed three times quickly in 2x SSCT and incubated for two hours in secondary antibody diluted in 2x SSCT. Biotinylated probes: sample was incubated in 1.5 μL of 488-conjugated streptavidin diluted in 98.5 μL detection solution (0.5 mL 1M Tris, 400 mg BSA, water to 10mL) for 1 hour, washed two times quickly in 2x SSCT, washed for 1 hour in 2x SSCT, and then washed 3 hours in 2x SSCT. (If using IDT-generated probes, these steps were not performed.) Ovarioles were mounted on a slide in 35 μL of DAPI + fluoromount.

Antibody for C(3)G(Anderson et al. 2005) was used. Figures 1b, 1c, 4d: Images of whole mount germaria were taken Zeiss LSM880 confocal laser scanning microscope using 63x/0.65 NA oil immersion objective with a 2x zoom using ZEN software. Figure 2b: Images were obtain using AIRY-Scan on Zeiss LSM880 confocal laser scanning microscope using40oil immersion objective. Images were saved as .czi files and processed using FIJI(Schindelin et al. 2012).

#### Live cell imaging

Ovaries were dissected in 10S Voltalef oil. The muscular sheath around each ovariole was removed and ovarioles were manually separated. Individual ovarioles were transferred to a drop of oil on coverslip. Videos were collected with an on an inverted Zeiss Axioobserver Z1 with motorized XYZ spinning-disc confocal microscope operated by Metamorph coupled to a sCMOS (Hamamatsuorca) camera and a temperature control chamber. All images were acquired with the Plan-Apochromat 100x/1.4 oil objective lens. Single-position videos in the germarium were acquired for 8 minutes at 25 ± 1 ° C, with a 10 second temporal resolution (12-slice Z-stack, 0.5 μm per slice).

### Quantification and statistical analysis

#### Recombination calculations

Genetic distances are expressed in centiMorgans (cM), calculated by 100 * (*R / n*), where *R* is the number of recombinant progeny in a given interval (including single, double, and triple crossovers), and *n* is the total number of progeny scored. 95% confident intervals were calculated from variance, as in Stevens (Stevens 1936). Molecular distances (in Mb) are from the positions of genetic markers on the *Drosophila melanogaster* reference genome, release 6.12 (Thurmond et al. 2018). Crossover density (or frequency), as calculated by cM/Mb, exclude transposable elements (see Miller *et al.*, 2016; Hatkevich and Sekelsky, 2017).

#### Quantitive microscopy analysis of IF in whole mounts

For fixed germaria, DAPI and anti-CID with either anti-C(3)G(Anderson et al. 2005) or anti-SMC1(Khetani and Bickel 2007) stains were used. Individual nuclei were first selected and eight 0.5 μm z-slices were used for analysis. First, fluorescence intensities were measured separately by subtracting cytoplasmic background in individual slices using an automated approach. For centromeric fluorescence intensities, centromeres were first segmented based on anti-CID (Active Motif) fluorescence using a probabilistic segmentation approach. Using segmented centromere masks, centromeric CID, C(3)G and SMC1 were quantified. For nuclear fluorescence intensities, nuclei were segmented using anti-C(3)G fluorescence. For chromosome arm fluorescence intensities, centromeric fluorescence intensities were subtracted from nuclear fluorescence intensities. To account for potential staining heterogeneity, fluorescence intensities were normalized to total nuclear CID fluorescence intensity, which was assumed to be unperturbed. Fluorescence intensities represent raw integrated densities of maximum intensity projected z-stacks. Nucleus selection, background subtraction and fluorescence intensity measurements were performed semi-automatically using custom FIJI-based plugins (available upon request) (Schindelin et al. 2012).

#### Analysis of pairing and centromere clustering

To determine the meiotic stage in fixed whole mount germaria, nuclear C(3)G staining patterning was used. Spots of C(3)G in early Region 2A was considered zygotene, full-length C(3)G in Region 2A was considered early pachytene, and full-length C(3)G in Region 3 was considered mid-pachytene. Two foci were considered unpaired if distances between the center of the foci were equal to or greater than 0.7 um (Gong, McKim, and Scott Hawley 2005). For centromere counting, any distinguishable single CID focus was counted as one, and distance between CID foci was not considered.

#### Live cell imaging tracking

The use of PAR1::GFP on live germaria allowed the identification of the different cyst stages. For live germaria, images shown are the projection of all Z-series of a single (t) projection. Three-dimensional tracking of spinning-disc data was performed using Imaris software (Bitplane). The CID::RFP signal was tracked using the ‘spots’ function with an expected diameter of 0.3 μm. Automatically generated tracks were then edited manually to eliminate inappropriate connections, including connections between foci in different nuclei or between foci of different sizes or intensity when more likely assignments were apparent or multiple spots assigned to the same focus.

To remove global movements of the germarium, each nucleus containing a CID::RFP focus was assigned to the nearest fusome foci. Then, the position of the reference fusome was subtracted from each CID::RFP focus for each time point of the tracking to get the relative tracks. These relative tracks were then compiled using a custom MATLAB (MathWorks) routine that computes the minimum volume of the ellipsoid that encloses all of the three-dimensional points of the trajectory.

To analyze centromere trajectories: Positions of individual centromeres were tracked every 10 seconds during 8 minutes to quantify the volume covered by each centromere. This raw volume was then corrected both for overall movements of the tissue and for variations in total nuclear volume. First, we subtracted the motion of the germarium using the position of the fusome as a reference within each cyst. Second, to take into account the nuclear volume at 8cc, we computed the relative volume, which is the raw volume divided by the mean value of the nuclear volume at 8cc stage. Finally, we normalized durations of each track by calculating the relative covered volume per second (as shown in Figure 3e).

#### Transparent Reporting

Each microscopy experiment performed in this study was repeated independently at least two times. We did not use explicit power analysis; rather, in each experiment, at least 8 independent germaria were imaged, and meiotic cells within the germaria were quantified, giving the final sample size per experiment. The total number of samples (n) is the sum of the final sample sizes per experiment.

## ACKNOWLEDGEMENTS

We thank the Sekelsky Lab, Scott Hawley, the Hawley Lab, and Abby Dernburg for critical review of this manuscript. We thank Sharon Bickel (SMC1 antibody and spreading protocol), Cathy Lake and Scott Hawley (C(3)G and Corolla antibodies), Kim McKim (*UAS::Smc1-HA* transgenic fly), Michaelyn Hartmann (IF/FISH protocol), and Bloomington Stock Center for generously providing reagents and protocols. Thanks to Tony Perdue and UNC Biology Microscopy Core for microscopy assistance. TH is supported in part by NIH grants 5T32GM007092 and 1F31AGO55157. Research in the laboratory of JS is supported by 1R35GM118127. VB is supported in part by predoctoral fellowships from the Fonds de Recherche Santé – Québec (FRQS). Research in the laboratory of PSM is William Burwell Harrison Fellow and supported by National Science Foundation CAREER Award 1652512. Research in the laboratory of JRH is supported by CNRS, Inserm, FRM(DEQ20160334884), ANR (AbCyStem), and Foundation Bettencourt-Schueller.

## AUTHOR CONTRIBUTIONS

Conceptualization, TH and JS; Methodology, TH, JS, VB, TR, and JRH; Investigation, TH, VB, and TR; Resources, JS, PSM, and JRH; Writing – Original, TH and JS; Writing – Editing, TH, JS, VB, TR, and JRH; Visualization, TH, VB, and TR; Supervision, JS, PSM, and JRH; Project Administration, TH; Funding Acquisition, TH and JS.

## SUPPLEMENTAL INFORMATION

Supplemental Information includes four figures, four tables, and six movies.

## SUPPLEMENTAL FIGURE LEGENDS

**Supplemental Figure 1 (related to Figure 2). Quantitative analysis of C(3)G in *WT* and *Mcm5^A7^* mutants**. a. Representative images of *WT* and *Mcm5^A7^* meiotic nuclei in whole mount germaria that were quantified in b., c., and Figure 2e examining C(3)G (magenta) and CID (green) in early pachytene. b. Quantification of C(3)G signal at the centromere (CID) in *WT* and *Mcm5^A7^* early pachytene nuclei. *p* = 0.4327, unpaired T-test. Data are represented as mean ± SD. c. Quantification of C(3)G signal at chromosome arm in *WT* and *Mcm5^A7^* early pachytene nuclei. *p* = 0.6358, unpaired T-test. Data are represented as mean ± SD. d. Representative images of *WT* and *Mcm5^A7^*meiotic nuclei of whole mount germaria that were quantified in e., f., and Figure 2f examining C(3)G (magenta) and CID (green) at mid-pachytene. e. Quantification of C(3)G signal at the centromere (CID) in *WT* and *Mcm5^A7^* mid-pachytene nuclei. *p* = 0.3615, unpaired T-test. Data are represented as mean ± SD. f. Quantification of C(3)G signal at chromosome arms in *WT* and *Mcm5^A7^* mid-pachytene nuclei. *p* = 0.5489, unpaired T-test. Data are represented as mean ± SD.

**Supplemental Figure 2 (related to Figure 4). Multiple germarium nuclei are depicted per frame**. (A, B, C) DAPI included images of *WT* and *Mcm5^A7^* in Figure 4a, b, c, respectively, to demonstrate that additional CID foci are of neighboring nuclei. d. DAPI included images of meiotic nuclei with 1 359-bp focus (WT, top panel) and 2 359-bp foci *(Mcm5^A7^*, bottom panel) from Figure 4d. Scale bars = 1 μm. Contrast and brightness of all images were adjusted for clarity.

**Supplemental Figure 3 (related to Figure 5). Images quantified in Figure 5**. a. Representative images quantified in Figure 5b and 5c. Scale bar = 1 μm. b. Representative images quantified in Figure 5d and 5e. Scale bar = 1 μm. Magenta: SMC1, Greed: CID. Images are of whole mount germaria.

**Supplemental Figure 4 (related to Figure 6). SMC1 overexpression in *Mcm5^A7^* mutants**. a. Representative images of *nos>Smc1, Mcm5^A7^* meiotic nuclei at meiotic onset, quantified in Figure 6a. b. Crossovers levels on Chromosome *X* in *WT (n*= 2179, 62.8cM (Hatkevich et al. 2017) and *Mcm5A^7^* (*n* = 2743, 3.8 cM), similar to levels previously reported(Lake et al. 2007). These data show that the crossover defect severity in *Mcm5^A7^* mutants is chromosome-specific. Due to genetics of the SMC1 transgene, we were unable to test *nos>Smc1; Mcm5^A7^* crossover levels on the *X*. Data are represented as mean ± 95% CI. See Table S4 for complete crossover dataset. c. Left: NDJ of the *X* chromosome in *WT* (0.07%, *n* = 3034) and controls *nos>Smc1; Mcm5^Df/+^* (0.16%, *n* = 1273) and *Mcm5>Mcm^WT^; Mcm5A^7^* (0.26%, *n* = 753). Right: NDJ of *rec^1/2^* (19.1%, *n* = 1563), and *nos>Smc1, rec^1/2^* (24.1%, *n* = 1187) to demonstrate that SMC1 overexpression NDJ rescue is specific to *Mcm5^A7^*. Data are represented as mean ± 95% CI.

**Supplemental Table 1 (related to Figure 2). Complete inter-sister recombination dataset**. Complete Ring:Rod dataset. *Adjusted females: Assuming male and female NDJ are equal, we subtract the amount of male NDJ from the *y* Normal female progeny; in these *y* females, one cannot distinguish between *y/y* female versus a *y/y/Y* female. *Ord^10^* exceptional females were distinguishable due to an additional phenotypic marker.

**Supplemental Table 2 (related to Figure 6). Recombination dataset for Chromosome 2L**. Recombination events across Chromosome 2L in progeny of *WT, Mcm5^A7^*, and *nos>Smc1, Mcm5^A7^* mothers.

**Supplemental Table 3 (related to Figure 6). Complete X-NDJ dataset**. Total normal and exceptional progeny from experimental and control lines.

**Supplemental Table 4 (related to Figure 6). Recombination dataset for *X* Chromosome**. Recombination events across *X* Chromosome in progeny of *WT, Mcm5^A7^*, and *nos>Smc1, Mcm5^A7^* mothers.

**Supplemental Video 1 (related to Figure 2). Rotation of *WT* meiotic nucleus shown in Figure 2b**. Meiotic nucleus demonstrating full length tracts of C(3)G (magenta) and localization of X-homologs (*X*-probes, green) in *WT*.

**Supplemental Video 2 (related to Figure 2). Rotation of *Mcm5^A7^* meiotic nucleus shown in Figure 2b**. Meiotic nucleus demonstrating full length tracts of C(3)G (magenta) and localization of X-homologs (*X*-probes, green) in *Mcm5^A7^*.

**Supplemental Video 3 (related to Figure 3). Dynamics of centromere clusters in 8-cell cyst nuclei in *WT***. Time lapse microscopy (spinning disc) expressing the centromere CID::RFP (magenta) and fusome marker Par-1::GFP (driven by the *nanos* promoter) (green). Frames were taken every 10 seconds.

**Supplemental Video 4 (related to Figure 3). Dynamics of centromere clusters in 8cell cyst nuclei in *Mcm5^A7^***. Time lapse microscopy (spinning disc) expressing the centromere CID::RFP (magenta) and fusome marker Par-1::GFP, (driven by the *nanos* promoter) (green). Frames were taken every 10 seconds.

**Supplemental Video 5 (related to Figure 3). Dynamics of one centromere cluster in one 8-cell cyst nucleus in *WT***. Time lapse microscopy (spinning disc) expressing the centromere CID::RFP (magenta) in one nucleus within an 8-cell cyst (dotted circle). Frames were taken every 10 seconds.

**Supplemental Video 6 (related to Figure 3). Dynamics of one centromere cluster in one 8-cell cyst nucleus in *Mcm5^A7^***. Time lapse microscopy (spinning disc) expressing the centromere CID::RFP (magenta) in one nucleus within an 8-cell cyst (dotted circle). Frames were taken every 10 seconds.

